# Influence of tropical pastures on grazing behavior parameters of young lambs

**DOI:** 10.1101/2020.11.09.374025

**Authors:** Jalise Fabíola Tontini, Cesar Henrique Espírito Candal Poli, Viviane da Silva Hampel, Mariana de Souza Farias, Neuza Maria Fajardo, Joseane Anjos da Silva, Luís Henrique Ebling Farinatti, James Pierre Muir

**Affiliations:** Department of Animal Science, Universidade Federal do Rio Grande do Sul, Porto Alegre, Rio Grande do Sul, Brazil; Department of Animal Science, Universidade Federal do Acre, Cruzeiro do Sul, Acre, Brazil; Texas A&M AgriLife Research, Texas A&M University, Stephenville, Texas, USA

## Abstract

Tropical sward characteristics can alter lamb ingestive behavior. Our study evaluated the ingestive behavior of young lambs in different tropical pastures to identify which variables interfere in their grazing activity. Two years of study were carried out with 54 weaned lambs distributed in three different swards: 1) monoculture of a upright grass, guinea grass (*Panicum maximum*; GG); 2) monoculture of a shrubby legume pigeon pea (*Cajanus cajan*; PP) and 3) contiguous areas with half the paddock with GG and half with PP (GP). The experiment was set out in a randomized complete block design. Lamb ingestive behavior was observed from sunrise to sunset with records every 5 minutes. To identify the main variables that affected with the lamb grazing activity, a multivariate analysis of the Decision Tree was performed. Our results showed that there was no difference in the ingestive behavior parameters of young lambs in different swards (*P* > 0.05). There was interaction among the swards and the experimental periods for the variables idleness time and biting rate (*P* ≤ 0.05). Grazing time of the animals increased 40% with experimental period progression. The Decision Tree identified leaf:stem ratio as the variable that most influenced lamb grazing time in GG and GP swards while in the PP sward grazing time was directly related to the pasture height. The behavior of young lambs on tropical pasture is variable as there is a change in the behavioral response over time. In addition, the grazing time of these animals can be estimated by means of variables related to pasture structural characteristics (leaf:stem ratio and height) together with chemical variables.

## Introduction

Knowledge of the plant-animal interface becomes an indispensable factor when working with pasture production systems. In pastoral ecosystems, herbivores develop grazing mechanisms or tools that make up what is called ingestive behavior [1]. Based on the existing literature, there are two main lines of research on ingestive behavior, one that explores animal response under different environmental conditions, with responses based on conditions extrinsic to animals [2–5], and the second that studies behavioral responses related directly to the different physiological conditions of animals, making a direct relationship between behavior and their intrinsic characteristics [6–8].

From an extrinsic perspective, the ingestive behavior may be one response, among other factors, to the type of pasture offered to the animal. In temperate pastures these behavioral responses are already well elucidated. There are conceptual models structured from scientific studies showing that pasture structure has an important effect on herbivore ingestive behavior characteristics and is directly responsible for the nutrient quantity ingested by grazing animals [3, 9–13]. However, studies of tropical pastures are scarce, and the existing results are inconclusive. In addition to the diverse growth habits and morphology of tropical forage species, there is structural variability over time.

In regions with tropical and subtropical climate, pastures are widely used in beef cattle production systems and appear to be expanding for sheep production. Despite this, the production of lambs on pastures with high productivity is challenging and some studies indicate unsatisfactory performance of the animals, below 100 g/day [14–16]. The productivity of lambs on pasture is attributed to their ability to harvest nutrients efficiently and effectively from the pasture [17], and the understanding of ingestive behavior is an important tool for directing management practices to obtain better animal performance. Therefore, before making adjustments to tropical pasture management to improve animal performance, it is essential to understand the dynamics of ingestive behavior of these animals.

In addition to the type of forage, the intrinsic aspects of the animals can also interfere in the ingestive behavior. Different physiological conditions of the animals require different nutritional inputs. In tropical grazing systems, sheep identify the heterogeneity of the physical and nutritional characteristics of plant components to meet their nutritional requirements. Diet selectivity is an animal strategy to avoid nutritional deficiencies and adaptive behavior allows grazing animals to perceive and interpret certain circumstances and thus modify their behavioral responses [18]. Assessing the behavior of grazing lambs on a fast-growing pasture is extremely important. In addition to the high nutritional demand, body size is associated with the time and energy spent selective grazing. The digestive capacity and body weight of these young animals are related to the choice of forage. This is one of the reasons why they have small mouths with the ability to select parts of a plant, generally resulting in a nutritional quality superior to the average offered in the pasture as a whole [19]. Therefore, to generate sustainable sheep production systems in tropical and subtropical environments, it is important to understand the factors that affect the ingestive behavior of grazing lambs [20, 21]. There is a lack of studies that show how lambs, which are highly selective and have important feed quality requirement and small mouth, graze upright tropical pastures with variable structure and nutritive value.

Many studies have already been carried out with the objective of understanding the ingestive behavior of animals grazing different species of forage [22, 23]. However, few studies have characterized the ingestive behavior of young lambs in different upright tropical pastures with the presence of grasses and/or legumes. Given the above, the objective of this work was to evaluate the ingestive behavior of lambs under conditions of continuous grazing submitted to different upright tropical species swards and to identify which variables influence the grazing activity of these animals.

## Material and methods

This study was approved by the Ethics Committee on Animal Use [Comissão de Ética no Uso de Animais da Universidade Federal do Rio Grande do Sul (CEUA-UFRGS) – project n° 27830] and conducted at the Research Station of the Universidade Federal do Rio Grande do Sul, Eldorado do Sul, Rio Grande do Sul State, Brazil (29 ° 13 ′26″ S, 53 ° 40 ′45″ W), 46 m above sea level. The climate is subtropical humid ‘cfa’ according to the Köppen (1948) classification. The cfa classification is characterized by hot summers with temperature averages over 22°C in the hottest month and rains distributed evenly over the year [24].

### Animals, treatments and experimental design

The experiment was repeated for 2 years (from 1 February to 25 April, 2015 and from 10 January to 12 April, 2016). The pasture and animal variables were evaluated every 21 days. The experimental area was 1.8 ha. Fifty-four castrated male lambs were used as testers, aged 3-4 months, weighing on average 22.6 ± 0.265 kg in Year 1 and 20.4 ± 0.310 kg of live weight (LW) in Year 2. Each year, 6 lambs were allocated per paddock, nine paddocks of 0.2 ha each were established, and these were considered experimental units. The animals were distributed by weight and fecal egg count (FEC), so that all paddocks had similar weight and total parasitic contamination of the animals. The animals were assigned to different tropical upright grass and legume swards: 1) monoculture of guinea grass (*Panicum maximum* cv. IZ-5; GG); 2) monoculture of pigeon pea (*Cajanus cajan* cv. anão; PP) and 3) contiguous areas with half the paddock with GG and half with PP (GP). The paddocks were arranged in a randomized block design. In order to evaluate lamb performance, they were weighed every 21 days, with prior fasting of liquids and solids for 12 hours.

### Pasture Assessments

The lambs remained on continuous grazing. Herbage allowance was adjusted every 21 days in order to provide same amount of leaf blade (ALB) from GG, PP and GP (kg DM/kg LW/ha/d) in the different paddocks, according to the methodology described by Sollenberger et al. (2005) [25]. The herbage allowance was adjusted to provide 10 kg DM ALB/100 kg LW/day to the animals in all paddocks, using the “put-and-take” technique [26]. According to this technique, there were two groups of animals, one called “testers”, represented by the animals that remain throughout the experimental period in grazing and express the effects of treatments, and the other group called “regulators”, used only to control the growth of the pasture and keep the herbage allowance at the desired level.

The height of the forage was obtained every 21 days. Fifty points were randomly measured in each paddock [27]. To estimate the herbage mass, six samples of 0.25 -m^2^ were cut close to the ground in each paddock, three of these represented the average height of the paddock and three were collected at random. Sub-samples of approximately 100 g were taken from each sample to carry out structural separation in leaf and stem plus sheath.

To evaluate the nutritional values of each feeding system, samples were collected every 21 days by grazing simulation, “hand plucking” technique [28]. These samples were dried in a forced-air oven at 55°C until constant weight and ground through a 2-mm sieve. This material was used to determine DM and crude protein (CP) according to the Association of Official Analytical Chemists [29], ether extract (EE) [30], neutral detergent fiber (NDF) [31] and acid detergent fiber (ADF) [32]. Total digestible nutrients (TDN) were obtained by the equation described by Sniffen et al. [33].

### Ingestive Behavior

Three evaluations of animal daytime ingestive behavior were performed [34]. The animals were observed from sunrise to sunset, recording every 5 minutes lamb grazing, ruminating or idling behaviour. In the interval between observations, biting rate was recorded for 20-bite time [35]. Within each paddock, the animal had different colored necklaces to facilitate identification.

### Statistical analysis

Analysis of variance was performed with repeated measures over time to determine the effects of treatments on the assessed variables of pasture and animals using the MIXED procedure of the statistical program SAS (version 9.4 - SAS Institute Inc., Cary, NC, USA). Treatment, block (within each year) and period were considered as fixed effects. Year and Treatment*block (within each year) interaction were considered as random effects.

The data were submitted to the Shapiro-Wilk normality test. Exponential transformation was performed on lamb average daily gain (ADG) data. Correlations among the variables were evaluated using Spearman correlation analysis.

The multivariate Decision Tree analysis was performed by JMP software (version 12 - SAS Institute Inc., Cary, NC, USA). This analysis indicates which were the factors that most affected animal response to grazing time. Independent variables included lamb initial body weight and biting rate as well as sward height, leaf:stem ratio, organic matter, crude protein, ether extract, mineral matter, neutral detergent fiber, acid detergent fiber and total digestible nutrients.

## Results

### Forage characteristics

Forage harvested by the animals in the different swards showed nutritionally adequate nutritional value, with crude protein values above 15% of DM and TDN close to 60% of DM. There was an interaction between sward type and period for all forage nutritive value variables (Table 1). The greatest amount of organic matter (*P* < 0.0001) was found in the last experimental period in the PP, and the smallest in the GG sward in the second period. The sward*period interaction for CP levels (*P* = 0.024) showed that GG had the lowest CP values, mainly in period 2 and 3. PP had the highest CP values, but it did not differ from the values found in the GP, independent of the evaluated period. The highest values of EE (*P* = 0.0412) were found in PP and in GP swards, mainly in the first and second period of the experiment. The values of mineral matter showed a significant interaction sward*period (*P* < 0.0001), PP presented the lowest values in the last evaluation period, while the GG presented the highest MM content in the final period of the study. The fiber content of the pasture (NDF and ADF; *P* < 0.0001) was higher in GG and GP swards and were even higher in the last evaluation periods. For TDN values (*P* < 0.0001), the interaction shows that, in swards with the presence of the grass (GG and GP), the amount of TDN decreased over the experimental periods. In PP, the decrease occurred only from the first to the second period, which shows that the legume can recover its nutritional value with the advance of production cycle.

**Table 1.**
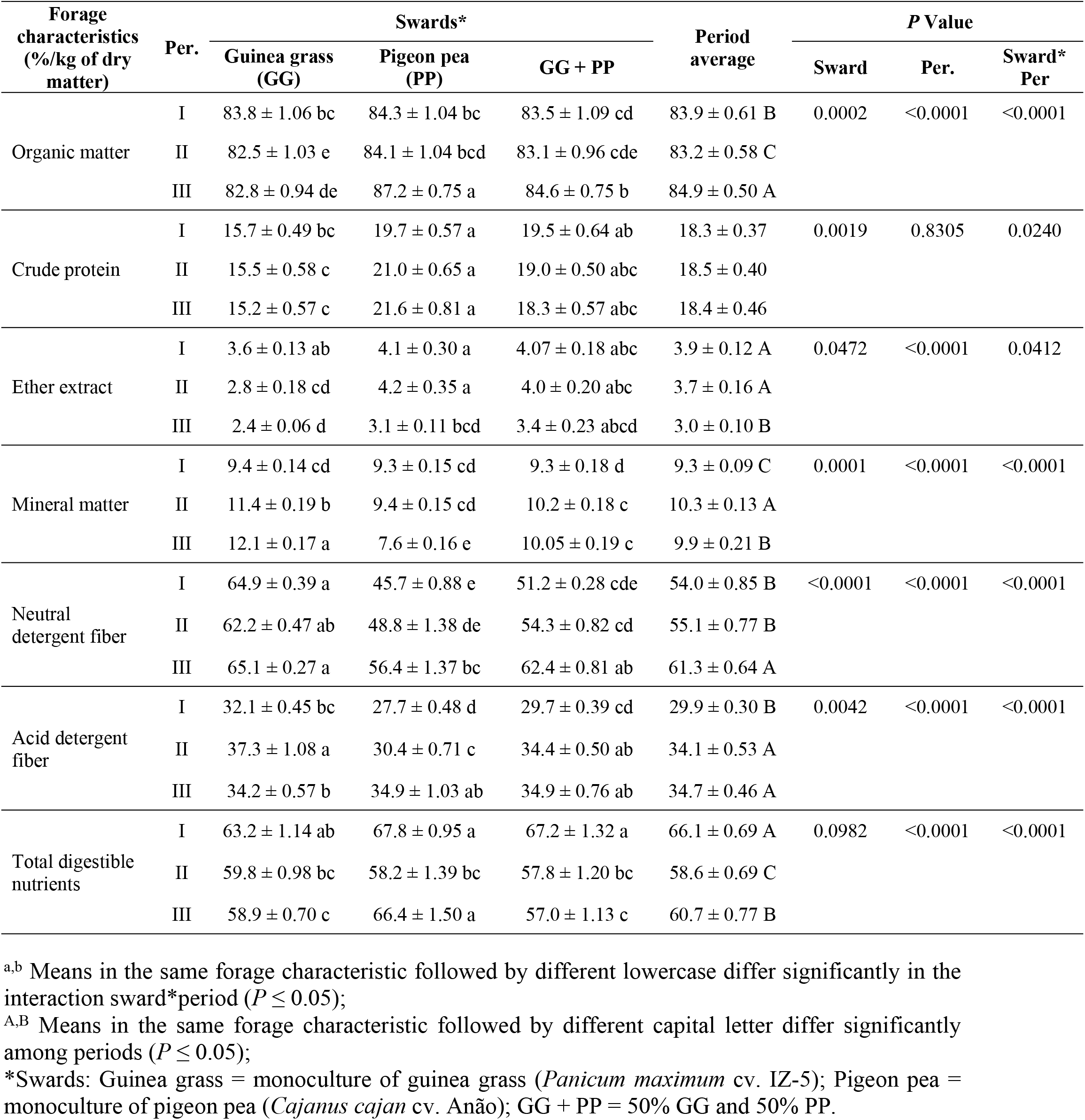
Qualitative characteristics of different tropical forages in the feeding of young lambs in southern Brazil. The means and their respective standard errors are presented.

| Forage characteristics<br>(%/kg of dry matter) | Per. | Swards* |  |  | Period average | P Value |  |  |
| --- | --- | --- | --- | --- | --- | --- | --- | --- |
|  |  | Guinea grass (GG) | Pigeon pea (PP) | GG + PP |  | Sward | Per. | Sward* Per |
| Organic matter | I | 83.8 ± 1.06 bc | 84.3 ± 1.04 bc | 83.5 ± 1.09 cd | 83.9 ± 0.61 B | 0.0002 | <0.0001 | <0.0001 |
|  | II | 82.5 ± 1.03 e | 84.1 ± 1.04 bcd | 83.1 ± 0.96 cde | 83.2 ± 0.58 C |  |  |  |
|  | III | 82.8 ± 0.94 de | 87.2 ± 0.75 a | 84.6 ± 0.75 b | 84.9 ± 0.50 A |  |  |  |
| Crude protein | I | 15.7 ± 0.49 bc | 19.7 ± 0.57 a | 19.5 ± 0.64 ab | 18.3 ± 0.37 | 0.0019 | 0.8305 | 0.0240 |
|  | II | 15.5 ± 0.58 c | 21.0 ± 0.65 a | 19.0 ± 0.50 abc | 18.5 ± 0.40 |  |  |  |
|  | III | 15.2 ± 0.57 c | 21.6 ± 0.81 a | 18.3 ± 0.57 abc | 18.4 ± 0.46 |  |  |  |
| Ether extract | I | 3.6 ± 0.13 ab | 4.1 ± 0.30 a | 4.07 ± 0.18 abc | 3.9 ± 0.12 A | 0.0472 | <0.0001 | 0.0412 |
|  | II | 2.8 ± 0.18 cd | 4.2 ± 0.35 a | 4.0 ± 0.20 abc | 3.7 ± 0.16 A |  |  |  |
|  | III | 2.4 ± 0.06 d | 3.1 ± 0.11 bcd | 3.4 ± 0.23 abcd | 3.0 ± 0.10 B |  |  |  |
| Mineral matter | I | 9.4 ± 0.14 cd | 9.3 ± 0.15 cd | 9.3 ± 0.18 d | 9.3 ± 0.09 C | 0.0001 | <0.0001 | <0.0001 |
|  | II | 11.4 ± 0.19 b | 9.4 ± 0.15 cd | 10.2 ± 0.18 c | 10.3 ± 0.13 A |  |  |  |
|  | III | 12.1 ± 0.17 a | 7.6 ± 0.16 e | 10.05 ± 0.19 c | 9.9 ± 0.21 B |  |  |  |
| Neutral detergent fiber | I | 64.9 ± 0.39 a | 45.7 ± 0.88 e | 51.2 ± 0.28 cde | 54.0 ± 0.85 B | <0.0001 | <0.0001 | <0.0001 |
|  | II | 62.2 ± 0.47 ab | 48.8 ± 1.38 de | 54.3 ± 0.82 cd | 55.1 ± 0.77 B |  |  |  |
|  | III | 65.1 ± 0.27 a | 56.4 ± 1.37 bc | 62.4 ± 0.81 ab | 61.3 ± 0.64 A |  |  |  |
| Acid detergent fiber | I | 32.1 ± 0.45 bc | 27.7 ± 0.48 d | 29.7 ± 0.39 cd | 29.9 ± 0.30 B | 0.0042 | <0.0001 | <0.0001 |
|  | II | 37.3 ± 1.08 a | 30.4 ± 0.71 c | 34.4 ± 0.50 ab | 34.1 ± 0.53 A |  |  |  |
|  | III | 34.2 ± 0.57 b | 34.9 ± 1.03 ab | 34.9 ± 0.76 ab | 34.7 ± 0.46 A |  |  |  |
| Total digestible nutrients | I | 63.2 ± 1.14 ab | 67.8 ± 0.95 a | 67.2 ± 1.32 a | 66.1 ± 0.69 A | 0.0982 | <0.0001 | <0.0001 |
|  | II | 59.8 ± 0.98 bc | 58.2 ± 1.39 bc | 57.8 ± 1.20 bc | 58.6 ± 0.69 C |  |  |  |
|  | III | 58.9 ± 0.70 c | 66.4 ± 1.50 a | 57.0 ± 1.13 c | 60.7 ± 0.77 B |  |  |  |

<sup>a,b</sup> Means in the same forage characteristic followed by different lowercase differ significantly in the interaction sward\*period ( $P \leq 0.05$ ); <sup>A,B</sup> Means in the same forage characteristic followed by different capital letter differ significantly among periods ( $P \leq 0.05$ ); \*Swards: Guinea grass = monoculture of guinea grass (*Panicum maximum* cv. IZ-5); Pigeon pea = monoculture of pigeon pea (*Cajanus cajan* cv. Anão); GG + PP = 50% GG and 50% PP.

Pasture productivity data showed that the different swards averaged >4000 kg DM/ha. With the use of the “put and take” technique, it was possible to maintain a similar green leaf allowance among the different swards in the different years (*P* > 0.05). Leaf:stem ratio (Table 2) showed no difference among swards with a mean of 0.51 (*P* = 0.1046). There were differences in leaf:stem ratio among periods, with a decrease over time (*P* < 0.0001). Pasture height was negatively correlated with leaf:stem ratio in PP swards (r = −0.60, *P* < 0.0001). Pasture height showed an interaction between treatment*period (*P* < 0.0001). The tallest height (134.2 ± 3.30 cm) was observed in the third period in PP, differing from other periods and swards (GG and GP). However, PP height in the initial and intermediate period of the study (103.2 ± 8.0 and 109.5 ± 6.9 cm, respectively) was not different from the height of GP (77.9 ± 2.43 cm), but it differed from the height found in GG (44.4 ± 1.50 cm). GP did not differ from GG, and the height of GG was different (*P* < 0.0001) among the different periods.

**Table 2.** Leaf: stem ratio of different tropical forages in the feeding of young lambs in southern Brazil at different experimental periods. The means and their respective standard errors are presented.

| Per | Swards* |  |  | Period average | P value |  |  |
| --- | --- | --- | --- | --- | --- | --- | --- |
|  | Guinea grass (GG) | Pigeon pea (PP) | GG + PP |  | Sward | Per | Sward*Per |
| 1 | $0.61 \pm 0.134$ | $0.50 \pm 0.149$ | $0.58 \pm 0.155$ | $0.56 \pm 0.022$ A | 0.1046 | <0.0001 | 0.0781 |
| 2 | $0.54 \pm 0.150$ | $0.38 \pm 0.113$ | $0.50 \pm 0.056$ | $0.47 \pm 0.022$ B | | | |
| 3 | $0.48 \pm 0.054$ | $0.46 \pm 0.055$ | $0.44 \pm 0.013$ | $0.41 \pm 0.025$ C | | | |

<sup>A,B</sup> Means followed by different capital letter differ between periods ( $P \leq 0.05$ );
\*Swards: Guinea grass = monoculture of guinea grass (*Panicum maximum* cv. IZ-5); Pigeon pea = monoculture of pigeon pea (*Cajanus cajan* cv. Anão); GG + PP = 50% of the paddock with guinea grass and 50% with pigeon pea.

### Animal performance and Ingestive behavior

Lamb ADG showed an interaction between sward and period (*P* < 0.0001), demonstrating that animals in GG and GP had greater capacity to maintain a constant ADG throughout the experimental periods than those grazing solely PP. The animals that grazed only legume (PP) had weight losses in the second period. Despite this interaction, there was no difference among treatments (*P* = 0.40), with mean values of 71 ± 8 for GG, 54 ± 9 for PP and 64 ± 8 g/day for GP. There was a correlation between leaf:stem ratio and ADG in PP (r = 0.42, *P* < 0.0001).

All variables of ingestive behavior were not only influenced by sward type, but also by period (*P* < 0.0001). The variations over time were important either independent or interacting with the sward type, as can be observed in Table 3. There was an interaction between sward type*period for idling time (*P* = 0.0390) and biting rate (*P* = 0.001). Idling time indicated that the longest times were recorded in the first period for all swards. For the biting rate variable, this interaction showed that, in the swards with the presence of grass (GG and GP), there was an increase in the number of bites per minute from the beginning to the end of the experiment. These values did not differ from the PP sward containing only the legume; however, in this case the biting rate did not change over the experimental periods.

**Table 3.** Means of ingestive behavior of lambs on tropical pasture in three different evaluation periods. The means and their respective standard errors are presented.

| Variables | Per. | Swards* |  |  | Period average | P Value |  |  |
| --- | --- | --- | --- | --- | --- | --- | --- | --- |
|  |  | Guinea grass (GG) | Pigeon pea (PP) | GG + PP |  | Sward | Per. | Sward *Per |
| Grazing (min.) | I | 360.4 ± 52.28 | 339.3 ± 58.79 | 367.4 ± 99.92 | 356.0 ± 74.11 C | 0.6100 | <0.0001 | 0.1964 |
|  | II | 450.9 ± 46.25 | 437.6 ± 60.77 | 455.5 ± 70.35 | 448.1 ± 59.96 B |  |  |  |
|  | III | 484.4 ± 114.25 | 499.8 ± 57.95 | 492.4 ± 89.80 | 492.2 ± 89.6 A |  |  |  |
| Idleness<br>(min.) | I | 248.7 ± 56.55 a | 251.8 ± 47.54 a | 226.9 ± 91.27a | 242.2 ± 68.49 A | 0.3287 | <0.0001 | 0.0390 |
|  | II | 126.2 ± 59.95 b | 148.2 ± 59.64 b | 122.3 ± 66.77 b | 132.2 ± 62.69 B |  |  |  |
|  | III | 131.76 ± 118.90 b | 139.6 ± 46.50 b | 135.4 ± 74.95 b | 135.6 ± 84.59 B |  |  |  |
| Rumination<br>(min.) | I | 110.9 ± 36.85 | 129.0 ± 54.59 | 125.7 ± 54.77 | 121.8 ± 49.61 B | 0.8814 | <0.0001 | 0.2163 |
|  | II | 143.2 ± 44.38 | 134.6 ± 36.89 | 132.3 ± 33.48 | 136.7 ± 38.38 A |  |  |  |
|  | III | 103.5 ± 38.21 | 77.6 ± 46.29 | 92.1 ± 41.93 | 91.1 ± 43.17 C |  |  |  |
| Bite rate<br>(per min) | I | 25.1 ± 11.05 cd | 25.3 ± 7.29 abc | 27.9 ± 8.81 bd | 26.1 ± 9.15 C | 0.9087 | <0.0001 | 0.001 |
|  | II | 27.6 ± 5.62 ab | 26.5 ± 9.39 abc | 25.7 ± 5.83 ac | 26.6 ± 7.1 B |  |  |  |
|  | III | 30.0 ± 5.32 ab | 25.8 ± 6.65 abc | 30.4 ± 7.1 a | 28.8 ± 6.69 A |  |  |  |

<sup>a,b</sup> Means in the same variable followed by different lowercase differ significantly in the interaction sward\*period ( $P \leq 0.05$ ); <sup>A,B</sup> Means in the same ingestive behavior followed by different capital letter differ significantly between periods ( $P \leq 0.05$ ); \*Swards: Guinea grass= monoculture of guinea grass (*Panicum maximum* cv. IZ-5); Pigeon pea= monoculture of pigeon pea (*Cajanus cajan* cv. Anão); GG + PP = 50% of the paddock with guinea grass and 50% with pigeon pea.

The grazing and ruminating times underwent changes in the different evaluation periods (Table 3). On average, there was an increase in the grazing time of approximately 40% throughout the experiment. The rumination time recorded in the second period was longer than the other periods. A positive correlation existed between the fiber content, represented by ADF, with grazing time (r = 0.55, *P* < 0.0001).

### Decision Tree analysis

Decision Tree analysis was performed with the grazing time data to understand which variables (already mentioned in the section “Statistical analysis”) influenced lamb grazing time in each sward. According to the analysis, in the GG paddocks, leaf:stem ratio and percentage of ADF of the forage explain 65% (R^2^ = 0.654) of the model, being the variables of greatest influence in animal grazing time. In PP paddocks, sward height along with TDN and ADF of the forage explained 57% (R^2^ = 0.571) of the model for grazing time. In GP swards, the percentage of ADF and leaf:stem ratio explained 65% (R^2^ = 0.649) of the grazing time observed.

As observed in Fig. 1, the Decision Tree model for GG showed that the variable with the greatest interference in grazing time was leaf:stem ratio in the first division of the decision tree, explaining 42% of model. Therefore, this analysis estimates that in pastures with leaf:stem ratio greater than 0.37, the average daily grazing time of the animals will be 404.8 minutes. If the leaf:stem ratio is less than 0.37, the grazing time of the animals will be greater, with estimated average of 565.3 minutes/day. In the second division of the Decision Tree model for GG, considering only the swards with leaf:stem ratio greater or equal to 0.37, ADF content appears as the second variable with the greatest influence on grazing time. It estimates that if GG has ADF >30.5% of DM, grazing time will be approximately 420 minutes/day. If this ADF is lower than 30.5%, grazing time decreases to 314 minutes/day.

**Fig 1.**
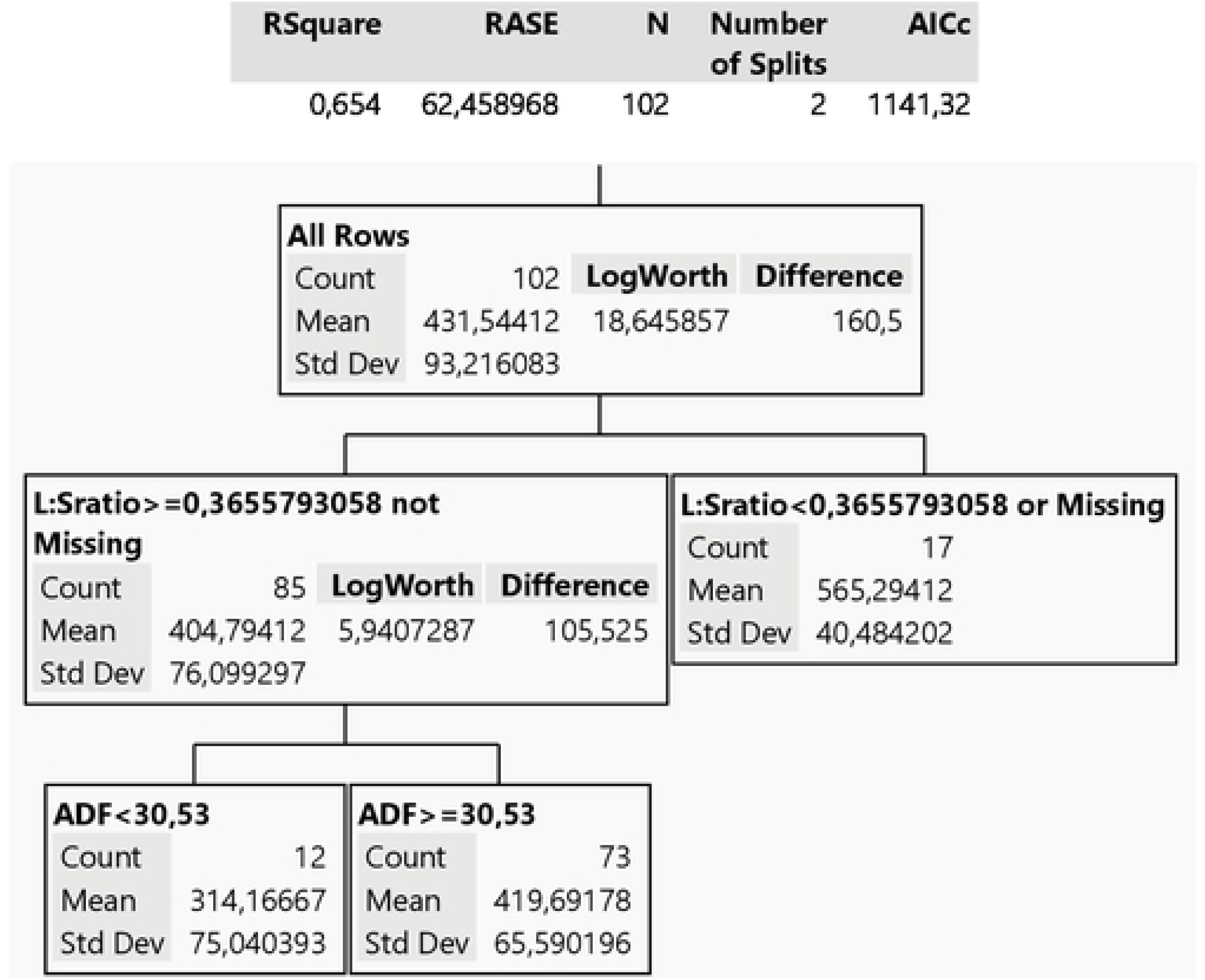
Decision Tree model for grazing time behavior variable of the animal grazing guinea grass (GG sward).

The Decision Tree model for grazing time in PP differed from GG (Fig 2). The main variable affecting lamb grazing time in PP is not related to the nutritional quality of the sward but, rather, its structure. The PP height contributed to 37% in the Decision Tree model. The model estimates that if PP height is < 76 cm, grazing time will be on average 329 minutes/day; however, if PP height is ≥76 cm, lamb grazing time will increase to 455 minutes/day. In the second division of the Decision Tree for PP, when height <76 cm, the percentage of TDN is the second variable that most affects grazing time. If TDN is greater than 60%, the estimated grazing time is 304 minutes/day. However, if that same pasture has a TDN <60%, the grazing time increases to 418 minutes/day. On the other hand, when PP is taller than 76 cm, ADF is an important determinant of grazing time. Therefore, in taller PP swards with ADF lower than 41%, lambs graze for 437 minutes/day, but when this same pasture has ADF >41%, the estimated grazing time is longer, 529 minutes/day.

**Fig 2.**
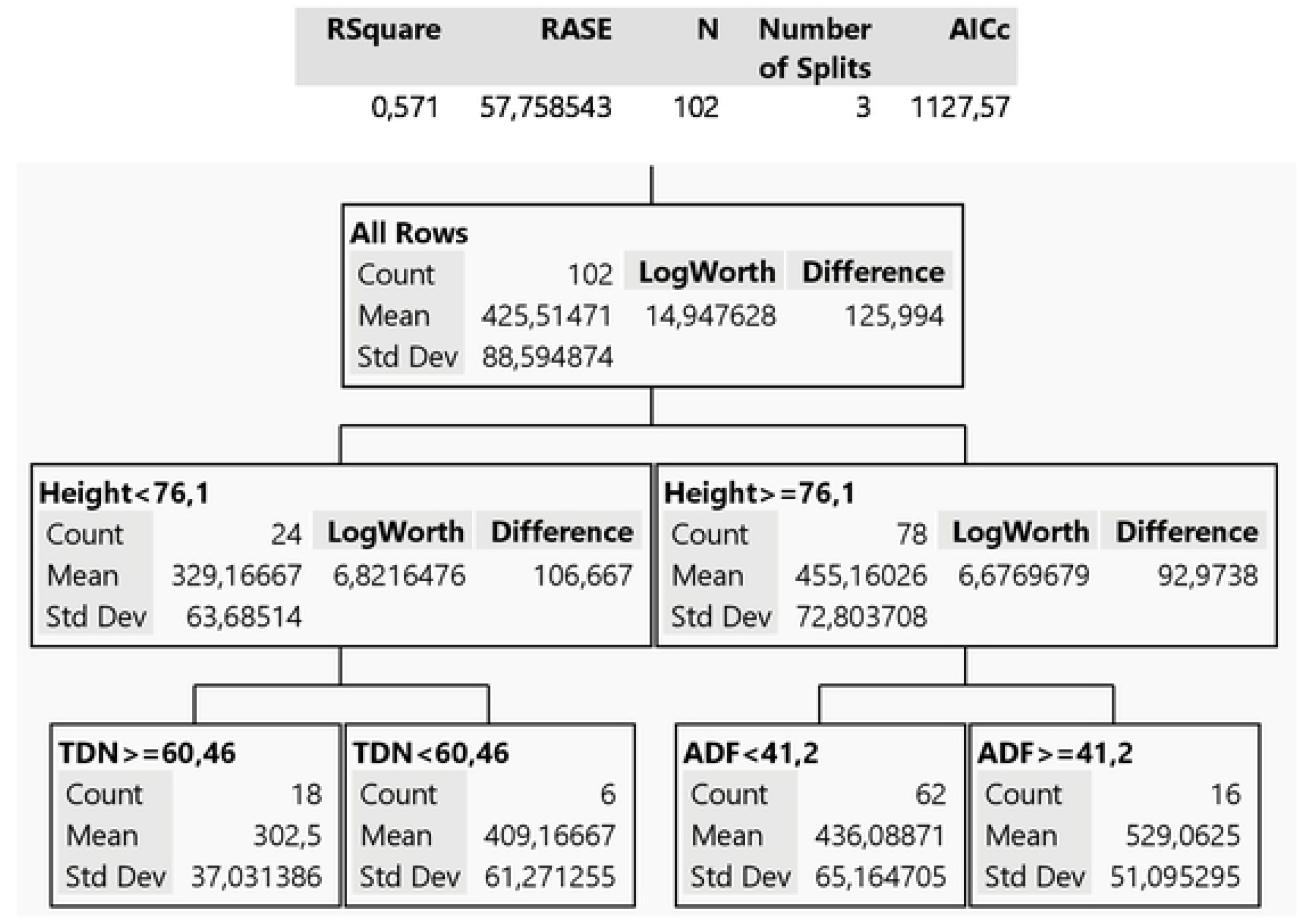
Decision Tree model for grazing time behavior variable of the animal grazing pigeon pea legume (PP sward).

For the GP paddock, the Decision Tree (Fig 3) showed that ADF content was the most important characteristic that determines grazing time, explaining 40% of the model. The mixed system of tropical upright grass and legume with ADF below 34% of DM allows an estimated grazing time 387 minutes/day, whereas when ADF content is >34% of DM, the estimated grazing time increases more than 25% (517 minutes/day). The second branch of the Decision Tree shows that, in addition to ADF, GP leaf:stem ratio can also explain the time that lambs spend grazing. In pastures with fiber content <34%, if the leaf:stem ratio is less than 0.63, grazing time goes from 387 minutes to 550 minutes/day, a 30% increase.

**Fig 3.**
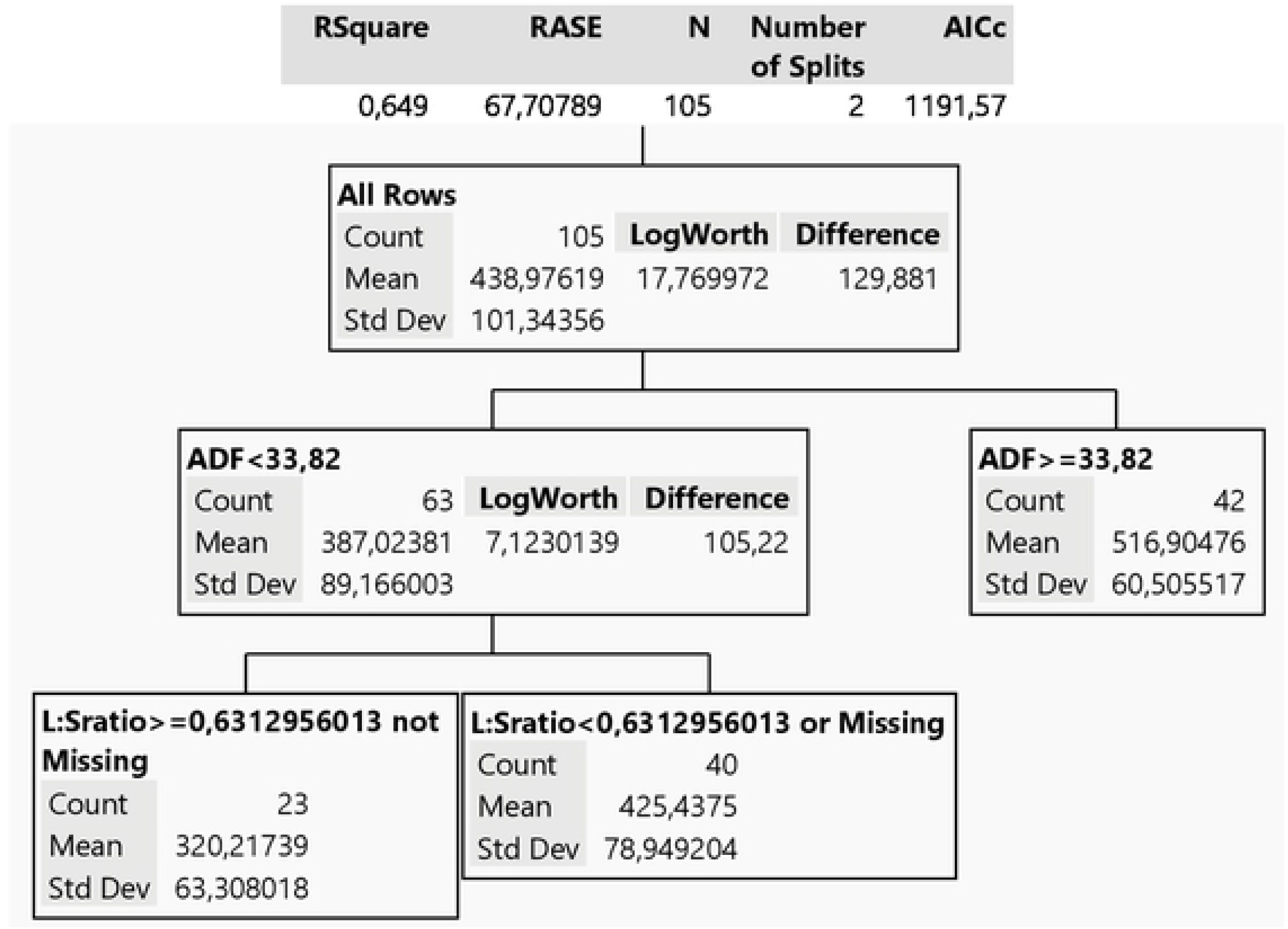
Decision Tree model for grazing time behavior variable of the animal grazing guinea grass + pigeon pea (GP sward).

## Discussion

We hypothesized that variable upright tropical pastures would affect ingestive behavior parameters of young lambs, as grasses and legumes present different structures e chemical compositions. Although, in our experiment, lambs had similar ingestive behavior in GG, PP and GP swards, when the data are analyzed over time, we observed an interaction among the different pastures and the experimental periods. These results indicated that changes in ingestive behavior occur and these are related to pasture morphological and histological and, consequently, nutritive values. Grazing time increased by 40% as experimental periods progressed. This change in grazing activity can be explained by the space-time variability that tropical pastures experience with the advancement of the production cycle, since pasture maturity influences forage structural characteristics and quality, with a decrease in new leaves and a concurrent increase in stem proportion of aboveground biomass [36–38]. Under these conditions, herbivores express behavioral adaptations to maintain the ingestion of a high-quality diet. Increasing grazing time can be an animal strategy to compensate for a reduction in grazing consumption in each bite [2, 22, 39]. High selectivity of lambs in searching for more nutritious plant parts such as leaves [40], may also require animals to graze longer because the proportion of leaves decreases over time (Table 2).

The decrease in quality with advancing stage of pasture maturity is well documented when analyzing the entire forage structure [36, 41–44]. Our data show that CP content did not change over time and energy values, represented by the TDN, but varied according to the sward. Although there was a decrease of TDN in PP, it increased from the second to the third period. In contrast, GG and GP showed a decrease in TDN content over time. The values found in this study are related to the nutritive value of the forage consumed by the animals and not by the entire plant structure, even if forage samples were collected by grazing simulation technique (“hand plucking”). This indicates that a decrease in nutritive value, with the maturation of the pasture as a whole, is not synonymous with lower quality of diet ingested by the animals. If lambs have the opportunity, they select the most nutritious plant parts, even with the low leaf:stem ratio observed in this study.

The quality and quantity of herbage mass available to the animals in this study did not limit lamb ADGs to <100 g/day [45–48]. The ADG in this trial were below expectations. According to the herbage mass and chemical composition offered to the animals in this study, NRC [48] estimates that lambs should achieve gains close to or greater than 100 g/day. These low ADGs were also observed with grazing lambs on guinea grass [14]. However, lambs grazing the tropical legume in our study with high productivity and nutritive value also showed lower-than-expected ADG. Lambs grazing exclusive legume pasture experienced a negative gain in the second experimental period. Our results indicate that the legume structure had a strong influence on low lamb performance once there were an increase of the legume height and a decrease of the leaf:stem ratio (r = −0.60, *P* < 0.0001). There was a significant correlation between leaf:stem ratio and ADG (r = 0.42, *P* < 0.0001). These results were also verified by another study [49] that suggested that, in tropical pastures, low animal performance is related to structural characteristics. The shrubby PP growth habit differs from forage legumes used in temperate pastures (*Trifolium* spp*., Medicago* spp.*; Vicia* spp., etc.). The characteristic PP woody stem, fast maturation rate and tall growth habit may make of this species a challenge for young lambs to graze.

Animals in the field use different strategies to increase or maintain consumption during grazing [2]. They can vary the mass and the frequency of the bite as well as grazing time. In our study, the increase in grazing time of the animals over the experimental period was not accompanied by the reduction of biting rate as described elsewhere [10]. Other studies, when evaluating an upright tropical grass, found that, when animals graze large leaves, they reduce bites per minute and graze for longer periods to achieve ideal leaf capture, manipulation, chewing and swallowing. In our study, we observed an opposite behaviour. Animals in the first period had less grazing time and lower bite rate but in the last period, the animals grazed for longer and obtained the greatest bite rates. Our results may be related to bite mass because, at the beginning of the study, there was a greater density of leaves (greater leaf:stem, Table 2). This plant structure allowed lambs to achieve greater bite mass and lower number of bites per minute. However, when leaf availability was more restricted and there was an increase in stem proportion, as happened in the second and third periods, bite mass was probably reduced and longer grazing time was necessary to maintain forage DM intake [15, 50, 51].

Rumination and idling time were relatively low in our experiment. Most ruminants spend more than 50% of the day resting and ruminating [52]. Another study [53] reported a rumination time of 466 minutes/day for lambs on millet pasture while our data showed an average of 120 minutes/12 hours of observation. However, rumination time was different among periods. In the last period of the experiment when plants were most mature, due to the greater time spent by the animals in the grazing activity, a reduction in rumination time occurred relative to that observed at the beginning of the study. Ruminating time may therefore be determined by the compensation between grazing and ruminating time during the day. Nevertheless, rumination time is also influenced by the nature of the diet and can be proportional to the cell wall content of forage [19]. In our study, there was no significant correlation between rumination time and forage cell wall content. We demonstrated that the time lambs spend ruminating in tropical upright pasture does not depend only on the pasture chemical characteristics, but also on pasture structure and other grazing activities. There is a trade off in time among grazing activities that makes grazing time a complex factor to be predicted [39]. Because of that, we carried out a Decision Tree multivariate analysis to determine the major factors that affect grazing time.

Multivariate analysis (Decision Tree) indicated that both physical and biochemical plant characteristics affect lamb grazing time. This result shows the major factors that affect a complex of drivers that determine lamb grazing time on tropical pasture. The vast majority of studies on animal behavior in pastoral systems report that grazing time is primarily related to the pasture structure, but there are studies that demonstrate that grazing time can be related to the pasture nutritional characteristics, mostly as a response of post-ingestive consequences influenced by previous feeding experiences [54].

The Decision Tree analysis shows that leaf:stem ratio is one of the main variables, related to the structure of tropical pastures, that influences grazing time in the GG monoculture and GP mixture. This result corroborates other studies [55, 56]. These authors comment that the way leaves are presented to the animals, and the degree to which grazing adaptations can avoid stems and less digestible dead materials, is of great importance in C4 pastures. The Decision Tree for GG estimated that a leaf:stem ratio >0.37 results in a 28% reduction in the grazing time of the animals when compared to a leaf:stem ratio <0.37. This reduction in grazing time was also reported in southern Brazil with *P. maximum* and *Brachiaria* spp. where grazing time decreased with increasing percentage of green leaves (r = −0.63 to −0.70) [57].

In addition to greater proportion of leaves allowing easier forage intake with better nutritive value, this characteristic becomes even more important with young grazing animals, as it allows a significant reduction in the time and energy spent on this activity. Reducing grazing time is directly related to the reduction of energy expenditure [58, 59], so animals can redirect energy for body development. Various physiological processes contribute to this relationship, such as the work of skeletal muscle for locomotion and the use of energy by the gastrointestinal tract and liver [60–62].

Another result that shows the usefulness of the Decision Tree analysis is the effect of forage ADF content on grazing time. Although the fiber content of the forage is usually related to rumination time [19, 63], there is a lack of knowledge relating the chemical characteristics of forages with grazing time. Therefore, this is an innovative result in describing ingestive behavior in tropical pastures. The ADF content showed a positive correlation with grazing time (r = 0.55, *P* < 0.0001). The higher the fiber content of the pasture, the greater the time spent by the animals grazing. As plant cell wall content increases, the proportion of leaves decrease, and the time spent by the animals to select more nutritious parts of the plants increases. The increase of grazing time is also associated to the need of the animals to maintain total DM intake during the day [39]. Bighorn sheep also show a negative relationship between grazing time and forage ADF content [64, 65], explained by ruminant behaviors to optimize nutrient extraction. As forage ADF content increases, the number of chews per bolus rises, increasing the manipulation for swallowing process and, consequently the time animals spend grazing.

A recent review [66] emphasizes that studies with tropical legumes are scarce regarding physical and chemical composition, and that these parameters are not considered as explanatory variables for animal responses. Therefore, the data from our study provide a clear relationship to the height of a shrubby tropical legume grazed by lambs. In particular, the PP, with a height of <76 cm may allow animals to spend less time grazing compared to when it is taller. Adjusting the legume pasture height, and indirectly its chemical composition (TDN and ADF), much as with grass swards, influence lamb grazing time. Tropical legumes have been used as an important source of protein in the production of ruminants [66, 67], however, ideal sward management for lamb production needs to be refined. Little is known about the correct way to make tropical legumes available to sheep in pastoral systems [67]. Our study indicates that grazing height of shrubby tropical legume, in our case PP, is an important factor for sheep grazing tropical pastures.

The Decision Tree analysis allowed us to visualize that, in tropical pastures that show rapid structural changes in a short period of time, it is possible to estimate the variation of sheep grazing time through relationships between forage plant physical and chemical pasture characteristics. Thus, our study is one of the first to report the influence of chemical and structural characteristics of tropical upright pastures on the time spent by lamb grazing animals, results that should guide future ruminant grazing management studies as well as practice.

## Conclusion

Our research shows that the behavior of grazing lambs in tropical upright pastures is variable, as there is a change in the grazing behavior over time, related to the structural and chemical modification of these pastures. Grazing time of these animals is strongly influence by structural (leaf:stem ratio and height) and chemical (ADF, TDN) characteristics of the pasture plants. Leaf:stem ratios in tropical pastures most influences lamb grazing time in upright perennial grass monoculture or in a mixed grass and legume swards, while, in shrubby legume monocultures, response in grazing time is directly related to pasture height. As such, management strategies that minimize shrubby legume height and stimulate leaf regrowth in both shrubby legumes and perennial bunchgrasses should enhance lamb performance on tropical pastures.

## Acknowledgments

This study was supported by grants from the National Council for Scientific and Technological Development (CNPq) of Brazil, and by Coordination for the Improvement of Higher Education Personnel (CAPES) of Brazil.

